# Accelerating joint species distribution modeling with Hmsc-HPC: A 1000x faster GPU deployment

**DOI:** 10.1101/2024.02.13.580046

**Authors:** Anis Ur Rahman, Gleb Tikhonov, Jari Oksanen, Tuomas Rossi, Otso Ovaskainen

## Abstract

Joint Species Distribution Modelling (JSDM) is a powerful and increasingly widely used statistical methodology in biodiversity modelling, enabling researchers to assess and predict the joint distribution of species across space and time. However, JSDM can be computationally intensive and even prohibitive, especially for large datasets and sophisticated model structures. To address computational limitations of JSDM, we expanded one widely used JSDM framework, Hmsc-R, by developing a Graphical Processing Unit (GPU) -compatible implementation of its model fitting algorithm. While our augmented framework retains the original user interface in R, its new computational core is coded in Python and dominantly uses TensorFlow library. This enhancement primarily targets to enable leveraging high-performance computing resources effectively, though it also accelerates model fitting with consumer-level machines. This upgrade is designed to leverage high-performance computing resources more effectively. We evaluated the performance of the proposed implementation across diverse model configurations and dataset sizes. Our results indicate significant model fitting speed-up compared to the existing Hmsc-R package across most models. Notably, for the largest datasets, we achieved *>*1000 times speed-ups. This GPU-compatible enhancement boosts the scalability of Hmsc-R package by several orders of magnitude, reaching a significantly higher level. It opens promising opportunities for modeling extensive and intricate datasets, enabling better-informed conservation strategies, environmental management, and climate change adaptation planning.

**Author summary:** Our study addresses the computational challenges associated with Joint Species Distribution Modelling (JSDM), a critical statistical methodology for understanding species distributions in biodiversity research. Despite its utility, JSDM often faces computational limitations, particularly for large datasets. To overcome this hurdle, we enhance the widely used Hmsc-R framework by introducing a GPU-compatible implementation of its model fitting algorithm. Our upgraded framework, while retaining the user-friendly R interface, leverages Python and TensorFlow for its computational core, enabling efficient utilization of high-performance computing resources. Through extensive evaluation across diverse model configurations and dataset sizes, we demonstrate substantial speed-ups compared to the original Hmsc-R package, with over 1000 times speed-ups observed for the largest datasets. This GPU-compatible enhancement significantly improves the scalability of JSDM, enabling the analysis of extensive and complex biodiversity datasets. Our work has far-reaching implications for informing conservation strategies, environmental management, and climate change adaptation planning by facilitating more efficient and accurate biodiversity modeling, ultimately contributing to better-informed decision-making in ecological research and practice.

## Introduction

The past decade has witnessed a transformative revolution in data acquisition and sampling methodologies, making large-scale data accessible for ecological research [1]. This emergence of novel data resources not only enhances our understanding of the biosphere but also establishes an innovative foundation for sustainable management, especially in the broader context of global change. Nonetheless, transmuting these extensive and intricate datasets into verifiable scientific insights poses significant challenges in terms of data processing and interpretation. The recently developed model-based approach, Joint Species Distribution Modelling (JSDM), presents a promising perspective, as it enable simultaneous evaluation of how environmental changes impact entire species communities [2, 3]. Several JSDM implementations have appeared as analytical software, exhibiting diversity in model structure assumptions and respective fitting algorithms [4]. In this context, we focus on the Hierarchical Modelling of Species Communities (HMSC) [5]. Notably, HMSC allows researchers to relate data on species communities to environmental predictors, species traits, and phylogenetic relationships. This regression-style modelling is conducted simultaneously with accounting for the study design characterising the data acquisition setting, which may include spatially or temporally explicit data [6]. The HMSC framework is implemented as an R package, Hmsc-R, that allows users to define and fit different models, and subsequently, post-process model outputs for prediction and inference [7]. Despite its utility, the increasing size of ecological datasets, coupled with the complexity of models designed for fitting these data, has often led to highly intensive computations in the existing implementation of HMSC [8]. This computational burden not only prolongs model fitting processes but also hinders the pace of scientific discovery and imposes limitations on ecological inquiries [9].

To address these computational bottlenecks, we developed Hmsc-HPC package that expands the Hmsc-R functionality by enabling efficient exploitation of Graphical Processing Units (GPUs) and High Performance Computing (HPC) infrastructure. The central innovation here lies in optimizing the model fitting process, which we shifted from being implemented via R-based calculations to TensorFlow-based computational backend that provides strategic utilization of GPU acceleration [10]. By leveraging the inherent parallel processing capabilities of GPUs, we expedite the execution of the block-Gibbs sampler used in HMSC fitting that consists of full-conditional updaters. This acceleration markedly reduces the time required for model fitting and, crucially, mitigates the performance bottleneck associated with input size; significantly increasing the capacity to tackle extensive datasets that were previously computationally prohibitive. Consequently, our technical enhancement facilitates a profound advancement in the domain of community ecology, paving the way for a deeper understanding of species-environment relationships and the dynamics of species communities within the context of larger and more intricate ecological systems, and potentially empowering decision-makers with critical insights for effective conservation and adaptation strategies in the face of rapidly changing ecosystems.

### Integrating Hmsc-HPC with Hmsc-R

Hmsc-R is an R package for joint outcome analysis, specifically tailored for ecologists studying species communities [7]. It encompasses a comprehensive statistical framework for integrating various commonly encountered ecological data types, including species occurrence records, environmental covariates, intricate study designs, species traits and phylogenetic information, to model species distributions and community assemblages [6]. From the statistical perspective, HMSC combines elements of generalized regression and latent factor approaches within a hierarchical modeling framework. The resulted statistical model is fitted in Bayesian paradigm via Markov Chain Monte Carlo (MCMC) sampling. This versatility allows it to account for a wide range of complex ecological processes and potential sources of variation and uncertainty. The Hmsc-R package features a user interface that enables ecologists to define models, fit models, and to utilize the fitted models for prediction and inference (Fig. 1):

**Fig 1.**
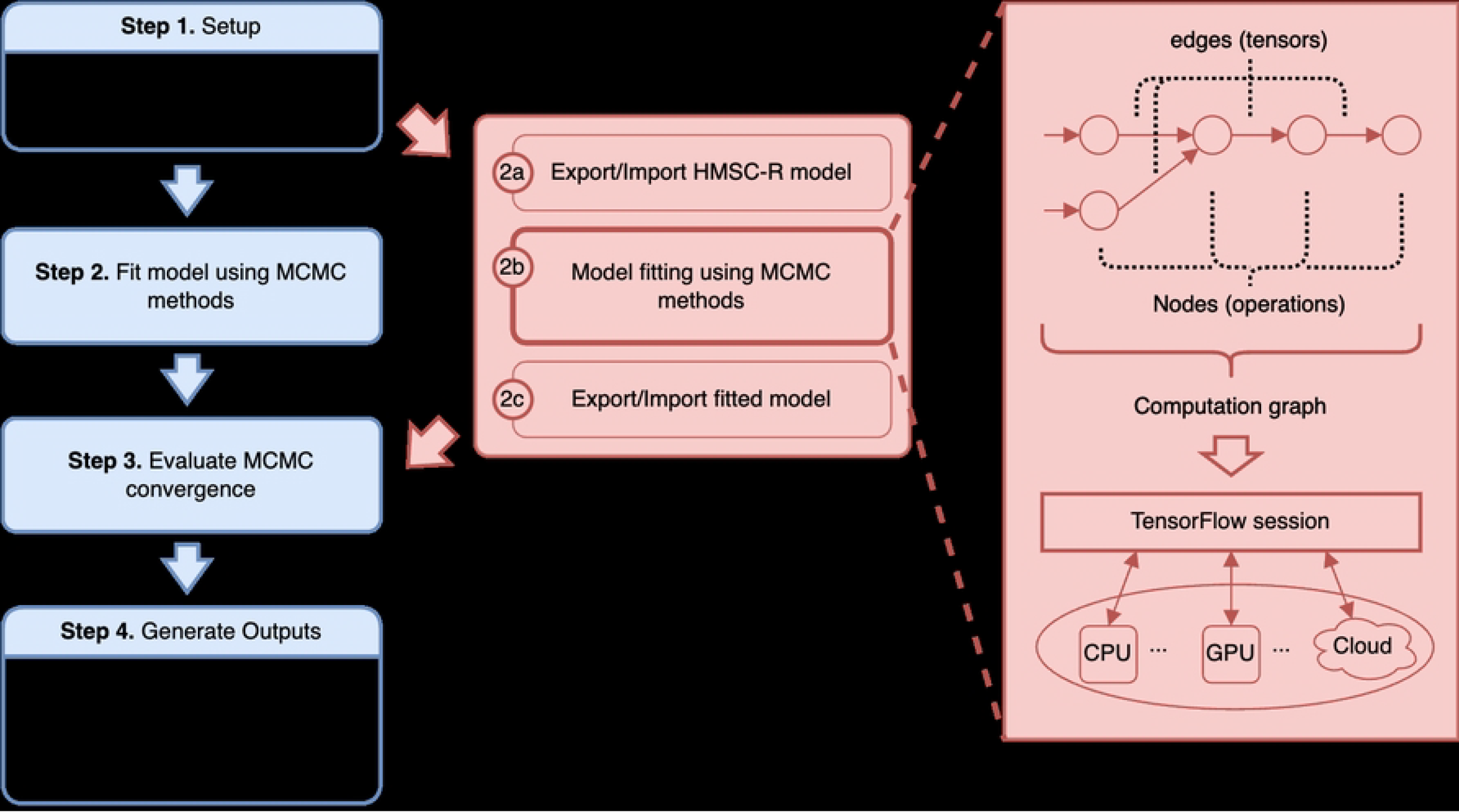
Pipeline of community analysis with HMSC. Left path represents the standard approach with Hmsc-R package [7]. The alternative on the right elaborates the newly-developed Hmsc-HPC augmentation that enables deployment of model fitting on HPC infrastructure.

#### 1. Setting model structure and data

In the first step the user defines the desired model structure, reflecting the specifics of the collected data and tailored to address specific ecological questions of interest. This step involves also pre-processing the original data to the assumed input format, specifying the set of optional model features, and, if necessary, fine-tuning model’s prior distributions.

#### 2. Model fitting

The defined model is then fitted to the provided data using MCMC sampling. Specifically, Hmsc-R utilizes a block-Gibbs sampler, which iterates over blocks (naturally defined subsets) of HMSC parameters. Within each block, parameters are collectively updated, conditional on all others being fixed to their current values, by sampling them from analytically derived conditional distributions. Due to the conditionally conjugate design of HMSC, these conditional distributions are available in closed form. The resulting MCMC chains are then processed by trimming and thinning to remove the effects of initial chains’ positions and autocorrelation. This refinement produces an approximation of the true, non-tractable posterior parameter distribution in HMSC, using a finite number of samples.

#### 3. MCMC fit diagnostics

In this step, the validity of the acquired MCMC chains is assessed via both formal criteria (e.g. Gelman-Rubin diagnostics [11]) and visual inspection of trace plots.

#### 4. Inference and predictions

After the reliable posterior approximation is obtained, this step encapsulates the operations that the user desires to perform with the fitted model; for instance, evaluating the model’s explanatory and predictive powers, exploring parameter estimates, and making predictions (for illustrations, see e.g. [5, 6]).

In practice, however, the use of Hmsc-R for large models is impeded by the prohibitive computational intensity of Step 2. The reason for that is two-fold. Firstly, Bayesian inference with MCMC methods is inherently more demanding than maximum likelihood alternatives, because the former seeks to assess the entire posterior distribution, which is a significantly more complex task than pinpointing an extreme point via optimization tools. Secondly, from a software engineering perspective, Step 2 in Hmsc-R relies primarily on native R computational routines, which are numerically inefficient compared to equivalent implementations in compiled programming languages [12]. In addition to impeding user experience, this reliance on a less efficient computational framework has led to grossly inferior computational performance of Hmsc-R when compared to several recently introduced alternatives designed specifically for highly accelerated model fitting [9].

In this work, our goal is to address the computational limitations of Hmsc-R and explore to what extent it can be enhanced once we substitute original model fitting implementation with a statistically equivalent replacement that is dedicatedly designed for High Performance Computing (HPC) hardware. We implemented our new package Hmsc-HPC, aiming for deployment to Graphical Processing Units (GPUs), which have proven to be effective for various computationally-intensive tasks beyond their original graphical applications over the recent decade. Despite the predominantly sequential nature of MCMC fitting algorithms, the computations within each MCMC step can significantly benefit from being broken down into small, independent tasks that can be executed simultaneously on the numerous GPU cores. Specifically, the bulk of statistical calculations within the block-Gibbs algorithm of Hmsc-R decomposes to linear algebra routines, many of which can be formulated within a “single instruction, multiple data” paradigm. These computations can be parallelized across the multiple processing units of a GPU, allowing for the simultaneous execution of calculations [13].

For our GPU-compatible augmentation, we redesigned and re-implemented the block-Gibbs sampler in Python programming language using the TensorFlow library [10]. We chose this numerical backend over lower-level GPU programming options due to TensorFlow’s well-developed application programming interface that features an extensive collection of linear algebra and statistical routines. This choice significantly reduced the need for developing standard low-level numerical operations or relying on third-party tools. Additionally, TensorFlow’s computation graph is a fundamental concept that underpins the framework’s functionality and efficiency. It represents the entire computation algorithm as a directed graph, where nodes denote mathematical operations, and edges represent the data flow between these operations. This graph-based approach offers several advantages, including portability, optimization opportunities, and distributed computing capabilities. Importantly, it decouples the definition of a computation from its execution, enabling deferred execution known as lazy evaluation. This separation allows TensorFlow to optimize the computation graph, fuse operations together for efficiency, and even potentially distribute the non-sequential parts of computation across multiple devices for concurrent processing. Graph execution, on the other hand, refers to the process of actually running computations on the graph, typically within a session context. It involves the evaluation of specific nodes or operations in the graph to produce results. This clear distinction between graph construction and execution not only enhances TensorFlow’s performance but also facilitates flexibility and extensibility in building complex data processing pipelines [14].

In addition to developing the Python code with TensorFlow-based block-Gibbs sampler implementation, we made appropriate modifications to the Hmsc-R package, enabling seamless integration of high-performance computing resources during the model fitting phase (Fig. 1):

#### 1. Setting model structure and data

In the initial step, the user specifies the model within R, following the standard workflow of Hmsc-R. This step also initializes the starting positions of the MCMC chains.

#### 2. Exporting specified model to Python

At this stage, the user saves the model as an RDS file in the R session, containing the model specification and MCMC settings.

#### 3. Model fitting with TensorFlow

In this core step, the user reads the RDS file into an independent Python session. Model fitting takes place in Python using the GPU-compatible MCMC sampler implemented in TensorFlow. This process begins with the compilation of the TensorFlow computational graph, which is then used for execution. Conceptually, this TensorFlow graph execution can be conducted at any device with properly set Python and TensorFlow dependencies. For instance it can be placed on the CPU in the very same modeller’s laptop as the other steps, and by itself it can provide a significant speed-up in model fitting. However, our implementation is designed to particularly target the subclass of GPUs dedicated for scientific computations, aiming to achieve ultimate performance boost for computationally intensive problems.

#### 4. Exporting the posterior to R

Once the posterior is obtained, the user stores it as an RDS file in the Python session.

#### 5. Diagnostics, inference and visualization, and prediction

In this step the user reads the RDS file back into an R session. Post-fitting steps continue within the standard workflow of Hmsc-R, independent of the changes made to the framework used for model fitting.

Finally, atop of intra-chain low-level parallelization that exploits the numerous GPU cores for performance boost, we also developed an interface enabling to compute several MCMC chains simultaneously. Thus, a common HPC infrastructure, such as a computational cluster, typically supports simultaneous placement of multiple jobs by a given user, and then the internal job scheduler automatically designates these jobs to available computational nodes. Since MCMC chains do not require any communication with each other, this provides an ideal opportunity for a parallelization across chains, with each job computing only a single or just a few chains.

The outlined design accomplishes two central objectives: it retains the established user interface of Hmsc-R as an R package while enabling the utilization of HPC infrastructure and GPU acceleration during the most computationally intensive step. This integrated, cross-language workflow combines the strengths of both R, Python and TensorFlow to offer a streamlined and efficient methodology for studying complex species communities within extensive ecological datasets.

## Performance comparison

We conducted a performance assessment of the GPU-accelerated HMSC implementation for different models characterized by varying data dimensions — in terms of number of sampling units and number of species used in the analysis. Additionally, we explored diverse model structures, encompassing considerations such as inclusion of phylogeny, design of random levels and choice of spatial method. The Hmsc-R execution, used as a baseline reference, was carried out and timed on a CPU of a consumer-level laptop. The Hmsc-HPC implementation was run on NVIDIA Volta V100 GPUs located in the AI partition of the Puhti cluster operated by CSC – IT Center for Science, Finland.

We set up our computational performance experiment with the same case study that was earlier used to compare the numerical performances of alternative spatial methods in the Hmsc-R implementation [15]. This case study serves as an illustrative example of the challenges encountered when using the Hmsc-R package with large datasets, even for non-sophisticated models. This dataset consists of spatially-referenced recordings of plant presence and absence in South-West Australia, along with several associated environmental attributes. In total, after excluding most rare species (those with fewer than five occurrences), the dataset comprises entries for 622 distinct species recorded at 25,955 locations. Additionally, it includes a small set of binary traits quantified across all recorded species. Furthermore, we accompanied it with the information regarding species’ taxonomy based on GBIF Backbone taxonomy [16].

We conducted experiments closely following the same experimental design as the original study [15] and evaluated the model fitting performance for five distinct variants of HMSC models potentially suitable for this dataset. These model variants cover different approaches to handle the spatial context of the data: 1) a spatially-ignorant model, 2) a spatial model with full Gaussian process (GP) structure for latent factors, 3) a spatial model with Predictive GP approximation (PGP, with 55 predictive process knots distributed along uniform hexagonal grid spanning the study area), and 4) a spatial model with Nearest neighbour GP approximation (NNGP, with ten neighbours). All of these four models incorporated all available traits but they did not utilize the information on taxonomy. To examine how the inclusion of taxonomy influences computational performance, model variant 5) represented a spatially-ignorant model that accounted for taxonomical relationships. We note that we use here taxonomy as a proxy of phylogeny, and that this model variant enables the estimation of phylogenetic signal in species responses to environmental predictors [5]. We introduced variation in two key dimensions: the number of species (*n*_*s*_ = *{*40, 160, 622*}*) and the number of sites (*n*_*y*_ = *{*100, 200, 400, 800, 1600, 3200, 6400, 12800, 25955*}*). This allowed us to create diverse subdatasets, ranging from reasonably small data size to fairly large data. To facilitate comparison across varying data dimensions, we kept the number of latent factors at the fixed value of 10. For each subdataset, we attempted to fit each of the five HMSC variants with both Hmsc-R and Hmsc-HPC, but excluded extreme cases with full GP spatial model that were infeasible due to insufficient RAM/VRAM. We recorded the total fitting time for each model-dataset combination and quantified the execution time required for single cycle of the block-Gibbs sampler. While different models may necessitate varying numbers of samples to converge, our primary focus here is the relative comparison between alternative implementations of the same mathematical algorithm. Therefore, the comparison of computational effort per Gibbs cycle provides a meaningful representation of the observed differences.

The results presented in Figs. 2 clearly indicate the computational benefits of fitting HMSC in a high-end specialized GPU device. The speed-up of Hmsc-HPC over Hmsc-R becomes larger as the data dimensions increase and the total computation time grows.

**Fig 2.**
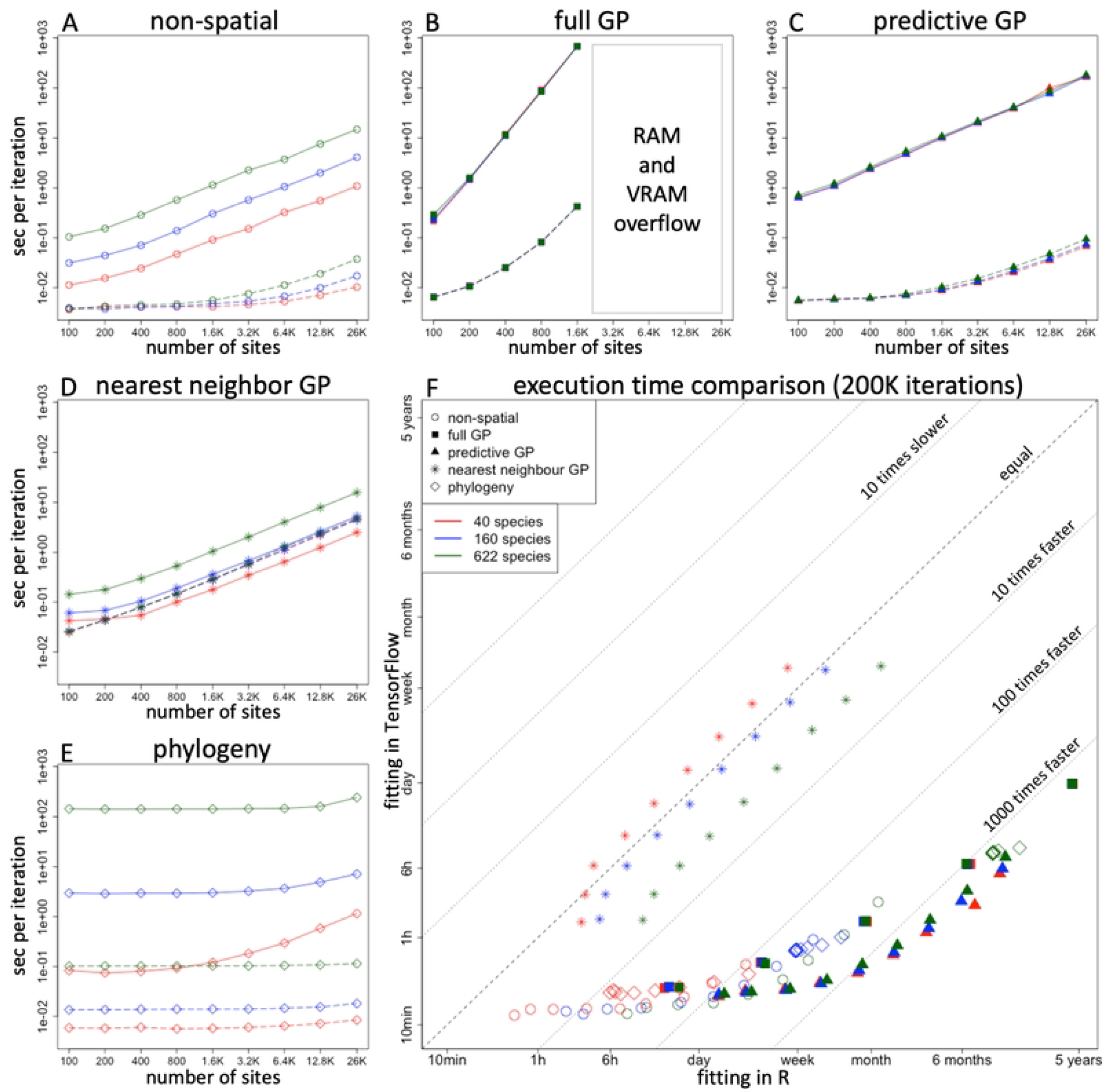
Performance comparison on GPU. Panels A-E represent execution time per block-Gibbs cycle for different models w.r.t. number of sites and number of species. Red, blue and green colors stand for *n*_*s*_ = 40, 160 and 622 species correspondingly. Solid lines depict the execution times for model fitting with R backend, and dashed lines for fitting with TensorFlow backend. In some panels lines of different colors overlap due to very minor effect of number of species on execution times. Panel F summarizes the overall comparison among execution times between R backend and TensorFlow backend. This figure shows the same data as depicted in panels A-E, but the times are multiplied by 200,000 to exemplify total computation times for a typical number of MCMC iterations in a model fit with transient of 100,000 and 1000 samples obtained with thinning 100. Dashed gray lines denote curves of designated ratios between TensorFlow and R computational performance (reads as TensorFlow is X times faster/slower than R). The symbols and colors refer to different model types and number of species, and they match the ones used in panels A-E (see legend).

Notably, even for a non-spatial model, in the case of the largest dataset, GPU-based execution outperforms the similar R-based execution by almost 1000 times, equivalent to an acceleration of 3 orders of magnitude. For full GP and PGP models, this acceleration becomes evident even with smaller numbers of sites, as these models involve operations with large matrices that are well-suited for GPU acceleration. The models that involve a taxonomy exhibit similar pattern, but aligned alongside the number of species rather than alongside variation in number of sites.

In contrast to the other model variants, the results on the NNGP models did not indicate apparent benefit of Hmsc-HPC use. HMSC algorithm for handling NNGP approximation for spatial latent factors relies on sparse linear algebra operations — namely Cholesky decomposition of sparse symmetric positive definite matrix and left-hand division with sparse triangular matrix. Presently, neither of these operations is available in TensorFlow. Hence, we relied on the capability to “inject” the required sparse operations from another Python package into TensorFlow graph, and consequently these operations are conducted on CPU. The amount of introduced device-host communication overhead appears to bottleneck the overall execution, diminishing the benefits of accelerated GPU execution for the rest of operations.

However, we note that as the methods developed here allow fast computation of GP and especially PGP model variants, we left solving the above mentioned bottleneck for future work as we considered a GPU-accelerated implementation of NNGP partially redundant.

## Conclusion

In this study, we introduced a novel Hmsc-HPC package that implements parallel and efficient implementation of HMSC fitting using GPU-compatible TensorFlow backend. This development is intentionally designed as essentially integratable add-on to the existing well-established Hmsc-R package in order to enhance its usability by JSDM practitioners. Our evaluation, conducted on an extensive dataset of species occurrence records, demonstrates that our implementation yields speed improvements of over 1000 times compared to the sequential R approach once the model fitting problem is sufficiently computationally intense (Fig. 2). This means that models that would earlier have required five years to fit can now be equivalently processed in one day, pushing significantly the boundary of what kind of HMSC models are feasible to fit.

In future work, we plan to explore the potential of inter-chain distributed computing and mixed floating number precision to harness the distinct features of modern GPUs and further improve the efficiency of Hmsc-HPC towards its ultimate extreme. Also, since our implementation is grounded on TensorFlow computational platform that supports auto-differentiation, we have expedited access to gradient-based generic MCMC samplers, e.g. Hamiltonian Monte Carlo [17], that can be integrated with existing block-Gibbs fitting algorithm of HMSC. The resulted hybrid MCMC sampling strategy would complement the drawbacks of each its components, yielding universally superior convergence properties for minor extra computational cost [18]. Finally, we aim to extend the evaluation of our implementation across a broader collection of diverse datasets and hardware configurations.

## Acknowledgements

We thank CSC – IT Center for Science, Finland for providing access to HPC infrastructure and high-end GPU devices. Matt White kindly provided a taxonomy of the plant species used in the case study. Graham Taylor and Sara El-Shawa are thanked for discussions and initial explorations related to alternative HPC software platforms among which we selected TensorFlow for this work.

This project was funded by the Academy of Finland (grant no. 336212 and 345110), and the European Union: the European Research Council (ERC) under the European Union’s Horizon 2020 research and innovation programme (grant agreement No 856506; ERC-synergy project LIFEPLAN), and the HORIZON-INFRA-2021-TECH-01 project 101057437 (Biodiversity Digital Twin for Advanced Modelling, Simulation and Prediction Capabilities).

## Authors’ Contributions

G.T. conceived the original core idea; G.T., A.R. and T.R. designed and developed the Hmsc-HPC implementation; J.O., A.R. and G.T. designed and implemented modifications for Hmsc-R; O.O. and G.T. designed the numerical experiments. All co-authors participated in discussions and contributed to paper writing.

## Data Availability Statement

The Hmsc-HPC add-on package is available for download from its designated GitHub webpage https://github.com/aniskhan25/hmsc-hpc. The repository includes preprocessed Australian plant data used in this study as CSV tables. Moreover, the repository also includes the R codes used in the Australian plant datas.

